# Why does the zebrafish *cloche* mutant develop lens cataract?

**DOI:** 10.1101/521526

**Authors:** Mason Posner, Matthew S. McDonald, Kelly L. Murray, Andor J. Kiss

**Author notes:** Corresponding author: Mason Posner 401 College Avenue, Ashland, OH 44805, USA.

## Abstract

The zebrafish has become a valuable model for examining ocular lens development, physiology and disease. The zebrafish *cloche* mutant, first described for its loss of hematopoiesis, also shows reduced eye and lens size, interruption in lens cell differentiation and a cataract likely caused by abnormal protein aggregation. To facilitate the use of the *cloche* mutant for studies on cataract development and prevention we characterized variation in the lens phenotype, quantified changes in gene expression by qRT-PCR and RNA-Seq and compared the ability of two promoters to drive expression of introduced proteins into the *cloche* lens. We found that the severity of *cloche* embryo lens cataract varied, while the decrease in lens diameter and retention of nuclei in differentiating lens fiber cells was constant. We found very low expression of both αB-crystallin genes (*cryaba* and *cryabb*) at 4 days post fertilization (dpf) by both qRT-PCR and RNA-Seq in *cloche, cloche* sibling and wildtype embryos and no significant difference in αA-crystallin (*cryaa*) expression. RNA-Seq analysis of 4 dpf embryos identified transcripts from 25,281 genes, with 1,329 showing statistically significantly different expression between *cloche* and wildtype samples. Downregulation of eight lens β- and γM-crystallin genes and 22 retinal related genes may reflect a general reduction in eye development and growth. Six stress response genes were upregulated. We did not find misregulation of any known components of lens development gene regulatory networks. These results suggest that the *cloche* lens cataract is not caused by loss of αA-crystallin or changes to lens gene regulatory networks. Instead, we propose that the cataract results from general physiological stress related to loss of hematopoiesis. Our finding that the zebrafish αA-crystallin promoter drove strong GFP expression in the *cloche* lens demonstrates its use as a tool for examining the effects of introduced proteins on lens crystallin aggregation and cataract prevention.

## Introduction

The ocular lens provides an excellent model system for examining tissue development and physiology due to its transparency, accessibility and the presence of only two cell types. Cataract, the development of opacities that interfere with the transmittance of light to the retina, continues to be the leading cause of human blindness worldwide [1]. A better understanding of the mechanisms leading to lens cataract could foster the development of preventative strategies. In recent years the zebrafish has become a powerful model for examining eye lens biology [2,3]. Not only do their transparent, external embryos facilitate experiments with the lens, but it is also relatively easy to express introduced proteins and explore their impact on lens function [4,5]. Multiple studies have shown that lens development and protein content are well conserved between zebrafish and mammals, making zebrafish studies translatable to our understanding of human lens disease [5-9].

Several zebrafish mutants have been described that exhibit lens cataracts during early development [10-12]. One of these, the *cloche* mutant, was first recognized by its cardiovascular system phenotype [13]. The homozygous *cloche* mutant lacks most endothelial and hematopoietic cells and does not survive past one-week post fertilization. A recent study identified a specific transcription factor gene affected in this mutant [14]. How this mutation leads to the lens phenotype is unclear. Goishi et al. [15] published the first study on the *cloche* lens, showing that mutant embryo lens fiber cells do not denucleate normally as they differentiate from the surrounding epithelial cell layer. They also showed that *cloche* lenses contained insoluble γ-crystallins, which may be the proximate cause of the lens opacity. Interestingly, the authors measured reduced expression of the lens small heat shock protein αA-crystallin in *cloche* lenses compared to wildtype embryos at 2, 3 and 4 days post fertilization (dpf) by relative end-point RT-PCR. Furthermore, they showed that overexpression of introduced αA-crystallin by mRNA injection could rescue the lens opacity phenotype. The authors concluded that the reduction in αA-crystallin led to γ-crystallin insolubility and resulting cataract in the *cloche* phenotype embryo.

While the conclusion that reduced expression of αA-crystallin leads to *cloche* lens cataract fits with our current understanding of the role α-crystallins play in maintaining lens transparency, there are several observations that suggest that other factors may be contributing to cataract formation in this zebrafish mutant. First, we have previously shown that suppressing αA-crystallin translation using synthetic RNA morpholinos does not produce a lens phenotype, even though αA-crystallin protein levels are reduced to undetectable levels by western blot [16,17]. Second, Zou et al. [18] showed lens abnormalities in a zebrafish αA-crystallin knockout line that are more subtle than the phenotype in the *cloche* lens with no reduction in lens size. And third, while microinjection of zebrafish αA-crystallin mRNA [15] and introduction of a rat αA-crystallin transgene [18] both reduced the severity of *cloche* lens cataract, it is possible that these proteins are hindering protein aggregation triggered by a mechanism other than loss of αA-crystallin.

The goal of this present study was to further characterize the *cloche* lens phenotype and revisit its possible causes to facilitate this mutant’s use as a model for studies on cataract development and prevention. We measured the levels of α-crystallin expression in *cloche* embryos by quantitative RT-PCR and conducted a global analysis of *cloche* transcriptomics by RNA-Seq. We describe variation in the severity of the *cloche* lens cataract and examine any correlations with changes in lens diameter and fiber cell differentiation. Lastly, we measured the abilities of a well-characterized human βB1-crystallin promoter and a native zebrafish αA-crystallin promoter to drive the expression of introduced protein into the *cloche* lens. In total, these experiments suggest that neither loss of αA-crystallin nor disruption in lens gene regulatory networks are the cause of the *cloche* lens cataract. We propose instead that the cataract results from a general physiological stress that triggers protein aggregation in the lens.

## Materials and Methods

### Fish maintenance

All protocols used in this study were approved by Ashland University’s Animal Use and Care Committee (approval # MP 2015-1). Wildtype ZDR strain and *cloche^m39^* adults were maintained on a recirculating filtering system at approximately 28°C with a 14:10 hour light and dark cycle. Fish were fed twice each day with live *Artemia* brine shrimp or flake food. Fish were bred at the most once per week to collect embryos for observation and microinjection. Adult *cloche^m39^* individuals were obtained from the Zon Laboratory at Harvard University and their genotype was confirmed by PCR using primer sets described by Reischauer et al. [14].

### Visualization of lens phenotypes

Embryos produced by incrossing *cloche* heterozygote fish were incubated at 28 °C in fish system water and transferred to 0.2 mM PTU at 6–30 h post fertilization to block melanin production. During microscopic visualization or collection for histology embryos were anesthetized in 0.016% tricaine. Lenses from whole live embryos were visualized by differential interference contrast microscopy on an Olympus IX71 inverted microscope and images were captured with a SPOT RT3 camera. The SPOT software was used to measure the diameter of each lens. Lenses were assigned to one of four classes based on the severity of the lens phenotype. Severity 3 lenses contained large central irregularities, severity 2 lenses contained small central irregularities and severity 1 lenses contained no central irregularity, but lacked the normal concentric lines formed by fiber cells in wildtype and sibling lenses. Embryonic lenses were also cryosectioned and stained with DAPI as described in Posner et al. [16]. The 4 dpf timepoint was selected for this analysis as fiber cell nuclei are typically removed in wildtype embryos by 3 dpf, allowing an additional day of development to gauge delay in the *cloche* embryos. We did not observe a noticeable qualitative change in the *cloche* cataract phenotype between 3 and 4 dpf. Embryos were euthanized by slow reduction of water temperature to freezing.

### Quantitative RT-PCR analysis of a-crystallin expression

Levels of αA-, αBa- and αBb-crystallin expression were measured in wildtype, *cloche* and *cloche* sibling embryos using qRT-PCR. Embryos were collected at 4 dpf and chilled on ice before replacing system water with RNAlater (Thermo Fisher) and then stored in a –20 °C freezer until RNA purification. Embryos were stored between 1 h and several days. Approximately 20 embryos were poooled for RNA purification from each genotype for each biological replicate. Three biological replicates were collected from independent fish crosses. Total RNA from each sample was purified using an RNEasy Minikit (QIAGEN) with Qiashreddor and quantified with a NanoDrop 1000 Spectrophotometer (Thermo Scientific). Purified total RNA (2,000 ng) from each sample was treated with DNasel (NEB) and 12 μl was used to synthesize cDNA using the Protoscript II First Strand cDNA Synthesis Kit (NEB) with the oligo d(T)23 primer in a total volume of 40 ul. The resulting cDNA sample was calculated to contain the equivalent of 16 ng/μl of original purified RNA.

All further procedures were identical to those described in Posner et al. [5] except that two reference genes were used instead of three (*ef1* and *rpl13a*). In short, three biological replicates for each genotype were amplified in technical triplicate using Luna Universal qPCR Master Mix (NEB) on an Applied Biosystems StepOne Real-Time PCR System (Thermo Fisher). Primer pair design for the two endogenous control genes and three zebrafish α-crystallin genes were previously published [19-21], and we previously validated the efficiency of these primers by standard curve and determined that they produced single amplification products [5]. Reaction conditions were identical to those previously published [5] and identical analysis was done to determine delta Cq values (using recommended MIQE guideline nomenclature [22]), which were then visualized and statistically analyzed using R [23] and R Studio [24].

### Comparison of *cloche* and wildtype gene expression by RNA-Seq

Between 10 to 20 embryos preserved in RNA*later* (Thermo Fisher) were removed from solution and homogenised in 200 μL of homogenisation solution in a Seal-Rite 1.5 ml microcentrifuge tube (USA Scientific) using a Microtip probe on a Fisherbrand^™^ Model 705 Sonic Dismembrator. Sonication was carried out for two periods of 10 seconds for a total of 10 Joules energy with a minute rest on wet-ice in between each disruption. Samples were judged to be homogenised when the sample was entirely homogeneous and no particulate matter settled to the bottom of the tube. Total RNA extraction was performed using a MAXWELL^®^ 16 LEV simplyRNA Total RNA Tissue Kit (Promega) as per manufacturer’s protocol and our previous reported methods [25]. Once isolated, total RNA was quantified by UV Nanodrop (Thermo Fisher NanoDrop 1000), and quality checked by Agilent BioAnalyzer2100 RNA Pico 6000 Chip Assay. Samples of total RNA with RINs below 8.0 were not used further.

Libraries for RNA-Seq compatible with Illumina short-read sequencing were constructed from the isolated, high-quality total RNA using the BIOO’s NEXTflex^™^ qRNA-Seq ^™^ Kit v2 using unique molecular indices (UMIs) [26-28]. The inclusion of RNA-Seq libraries that employ UMIs reduces PCR introduced bias during library construction, thus increasing the accuracy of the quantitative nature of differential gene expression (DGE) in downstream analysis. Library quality and quantification was validated using BioAnalyzer2100 HS DNA Chip Assay and KAPA Universal Illumina Library Quantification Kit. Three biological samples from each condition, wild-type (WT) and cloche phenotype (CP), were used to prepare libraries, for a total of six.

Libraries were quantitated, and frozen at -80°C and shipped on dry ice O/N by courier to the Center for Genome Research and Biocomputing (CGRB) at Oregon State University for sequencing on an Illumina HiSeq3000 platform. The six samples were loaded onto a single HiSeq3000 lane and data was acquired using a 2x150 bp paired-end run with a 10% PhiX spike-in to account for the low diversity of the UMIs within the first 9 nucleotides of both Read1 and Read2. Libraries were demultiplexed and raw FASTQ data retrieved from the CGRB (OSU) and processed at Miami University’s Center for Bioinformatics & Functional Genomics (CBFG).

Bioinformatics analysis was performed on CLC Genomics Workbench 11.0.1 on an AMD Opteron Workstation using 256 GB ECC RAM and a 12 TB storage RAID5 array running Ubuntu 16.04.5 LTS. Data were deconvoluted based on UMIs using the Molecular Indexing Plug-In (Toothfish Software) on CLC GW; trimmed, mapped to the zebrafish *Danio rerio* annotated genome, build GRCz10. Once reads were mapped, an RNA-Seq analysis experiment was performed (WT, CP). Statistical analysis was performed in a pairwise manner using the bootstrapped receiver operator characteristic (bROC) Plug-In 3.0 (BioFormatix, Inc). The use of the bROC Plug-In enables the non-parametric analysis of RNA-Seq data with low replicate numbers and enables a more robust and accurate DGE result [29,30]. The RNA-Seq experimental analysis was then annotated with GO term from within CLC GW and data exported to a CSV based spreadsheet for inspection. Data was visually plotted in CLC Genomics Workbench and plots were exported in SVG format for presentation in figures.

### Construction of expression plasmids, embryo microinjection and visualization of GFP expression

The construction of a plasmid driving expression of green fluorescent protein (GFP) using a 1kb zebrafish αA-crystallin promoter was previously described [5] and the plasmid is available from Addgene. A second plasmid driving GFP expression with a 296 bp fragment of the human βB1 crystallin promoter was obtained from the Hall laboratory at the University of California at Irvine, and has been previously reported to drive expression in zebrafish lens [31]. We compared the ability of both promoters to drive the expression of GFP in *cloche* embryos to determine the best promoter to use in future experiments testing effects of introduced proteins on *cloche* cataract. To prepare promoter expression plasmids for injection into zebrafish embryos, plasmids were linearized with *NotI* (NEB), purified with the Monarch PCR and DNA Cleanup kit (NEB), and then dialyzed with 0.5X TE buffer using a 0.025 μm VSWP membrane (Millipore, Billerica, MA, USA). Injection solutions contained 35 ng/ul of the dialyzed plasmid, 0.2% phenol red and 0.1 M KCl to bring the solution to 5 ul. Two nanoliters of this plasmid mix was injected into zebrafish zygotes with a Harvard Apparatus PL-90 picoinjector (Holliston, MA, USA) using needles prepared with a Sutter P97 Micropipette Puller (Novato, CA, USA). Injection pressures were adjusted to inject 1 nl of plasmid solution with each 20 ms pulse. Injected embryos were incubated at 28 °C in fish system water and transferred to 0.2 mM PTU at 6–30 h post fertilization to block melanin production and facilitate observation of GFP expression. Injected embryos were anesthetized in tricaine at 4 dpf and imaged at 200× total magnification using UV illumination and GFP filter on an Olympus IX71 inverted microscope. Images were captured with a SPOT RT3 camera (Diagnostic Instruments, Sterling Heights, MI, USA).

## Results

Homozygous *cloche* embryos were recognizable by the presence of cardiac edema (Fig 1A compared to 1B), an abnormally shaped heart (Fig 1C and D), bent trunks, reduced eye size compared to non-phenotypic *cloche* siblings, and the lack of circulating red blood cells. Some of these features, such as cardiac edema and bent trunks, are typical of many abnormal embryos phenotypes. However, the loss of red blood cells and reduced eye size was diagnostic for the mutation. We confirmed that our *cloche* fish heterozygote breeding population contained the *cloche^m39^* allele by PCR amplifying the identified region of mutation using primers from [14](Fig 1E).

**Figure 1.**
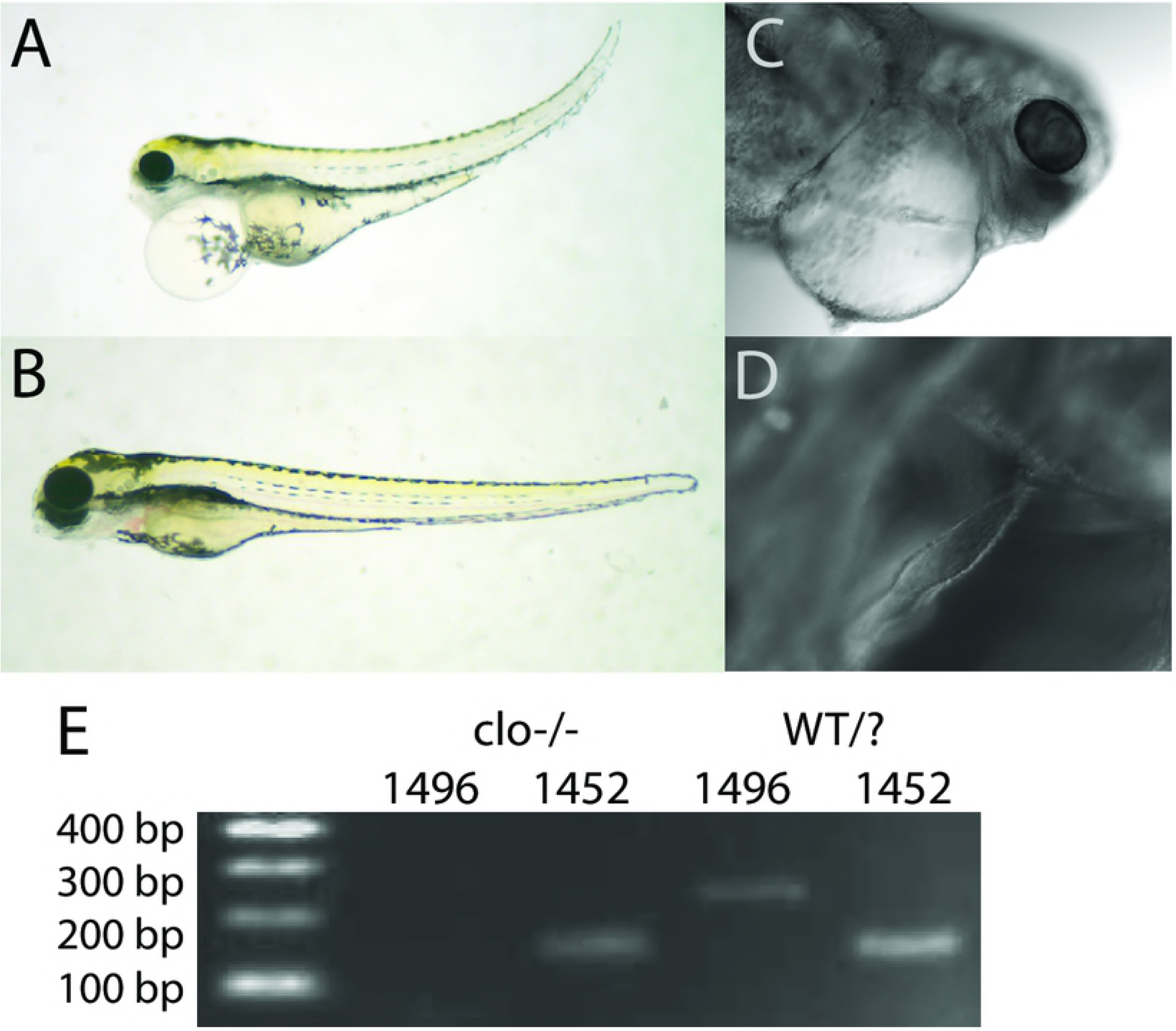
Identification of *cloche* embryos. View of the gross morphology of an embryo homozygous for the *cloche* mutant allele *m39* (A) compared to a non-phenotype sibling (B) at 4 dpf. Embryos were identified by the presence of cardiac edema, lack of red blood cells and characteristic irregularly shaped heart (C and D). The presence of the *m39 cloche* allele in our fish was confirmed by PCR genotyping using primer sets z1496 and z1452 (E; [14]).

### Severity of the *cloche* lens phenotype was variable

Previous work characterized abnormalities in the *cloche* lens and quantified light reflectance, retention of fiber cell nuclei and eye and lens size [15,32]. We noticed wide variation in the visible opacities that develop in *cloche* embryo lenses and quantified the range of this phenotype using a severity scale that placed each lens in one of four possible categories (Fig 2). Data from 4 dpf *cloche* lenses showed that 47% fell within the most severe category (Fig 2: Sev 3), but that 12% of lenses showed minimal disturbance in transparency (Fig 2: Sev 1). However, no *cloche* phenotype lens showed the normal, concentric rings found in sibling lenses (Fig 2: Sib). All observed *cloche* siblings, which would be a mix of heterozygotes and homozygous wildtype, had no noticeable abnormality in transparency.

**Figure 2.**
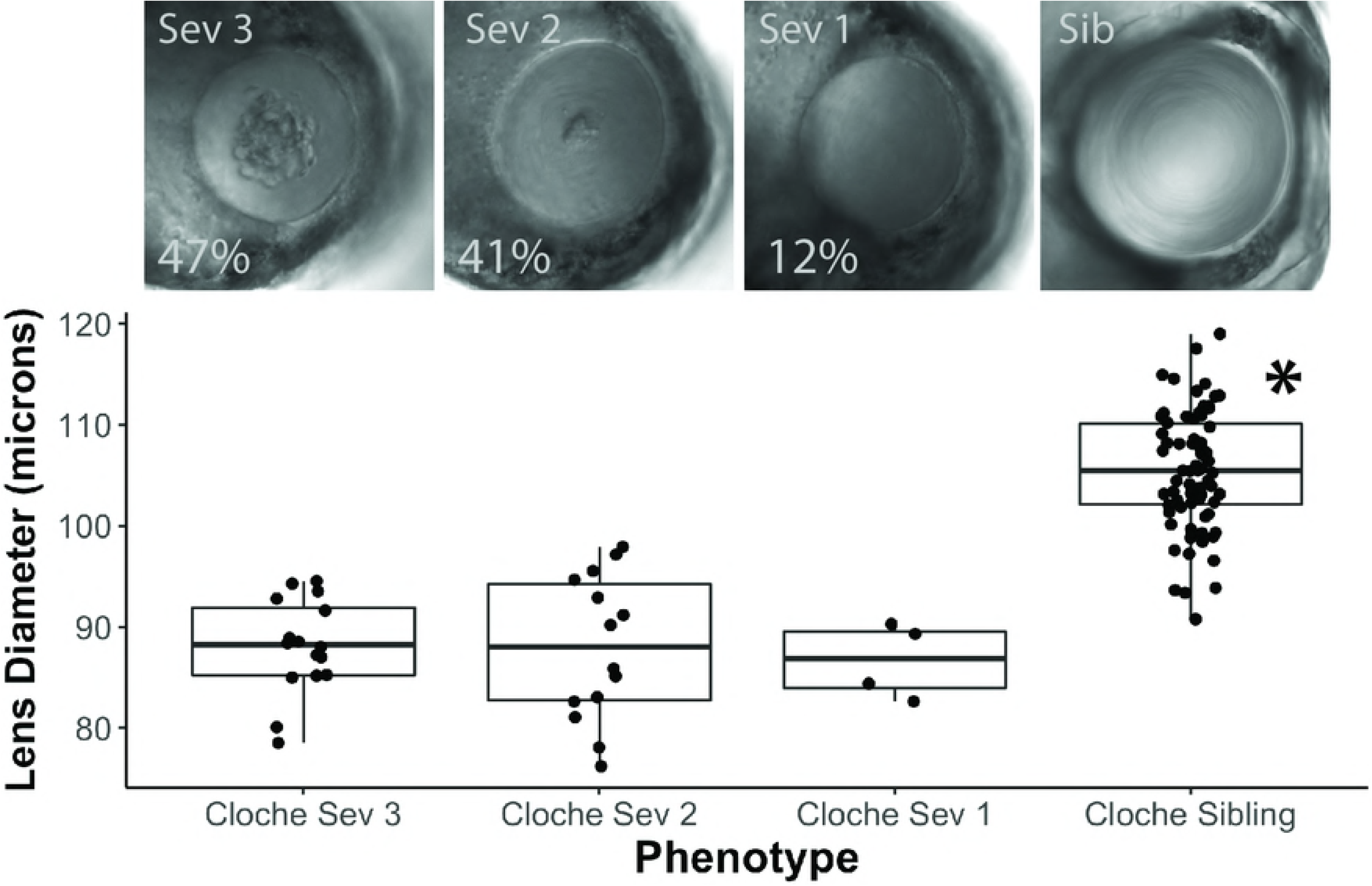
Severity of the cloche lens phenotype varies, but is not correlated with lens diameter. *Cloche* embryos at 4 dpf were pooled into three severity groups. Representative lenses are shown for severity groups 3, 2, and 1, with group 3 being most severe. Percentages indicate the proportion of embryos with each severity (n=34). A representative normal lens is shown from a *cloche* sibling. The lens diameter in *cloche* embryos was uniformly reduced in all severity groups compared to siblings (ANOVA *p* value < 0.0001; Tukey Honest Significant Difference (HSD) post test used to identify statistically significant mean for sibling group (*)).

We were interested in determining if smaller lens size and delay in fiber cell differentiation correlated with *cloche* lens opacity. Lens diameter of 4 dpf *cloche* embryos did not differ significantly between severity groups but were uniformly reduced compared to *cloche* siblings (Fig 2). We also found that fiber cell nuclei were retained in *cloche* embryos of all severities in comparison to *cloche* siblings, which retained no fiber cell nuclei at 4 dpf (Fig 3). We did see an unexpected statistically significant smaller average number of fiber cell nuclei in *cloche* embryos of the greatest severity (Fig 3: Severity 3 compared to 2 and 1).

**Figure 3.**
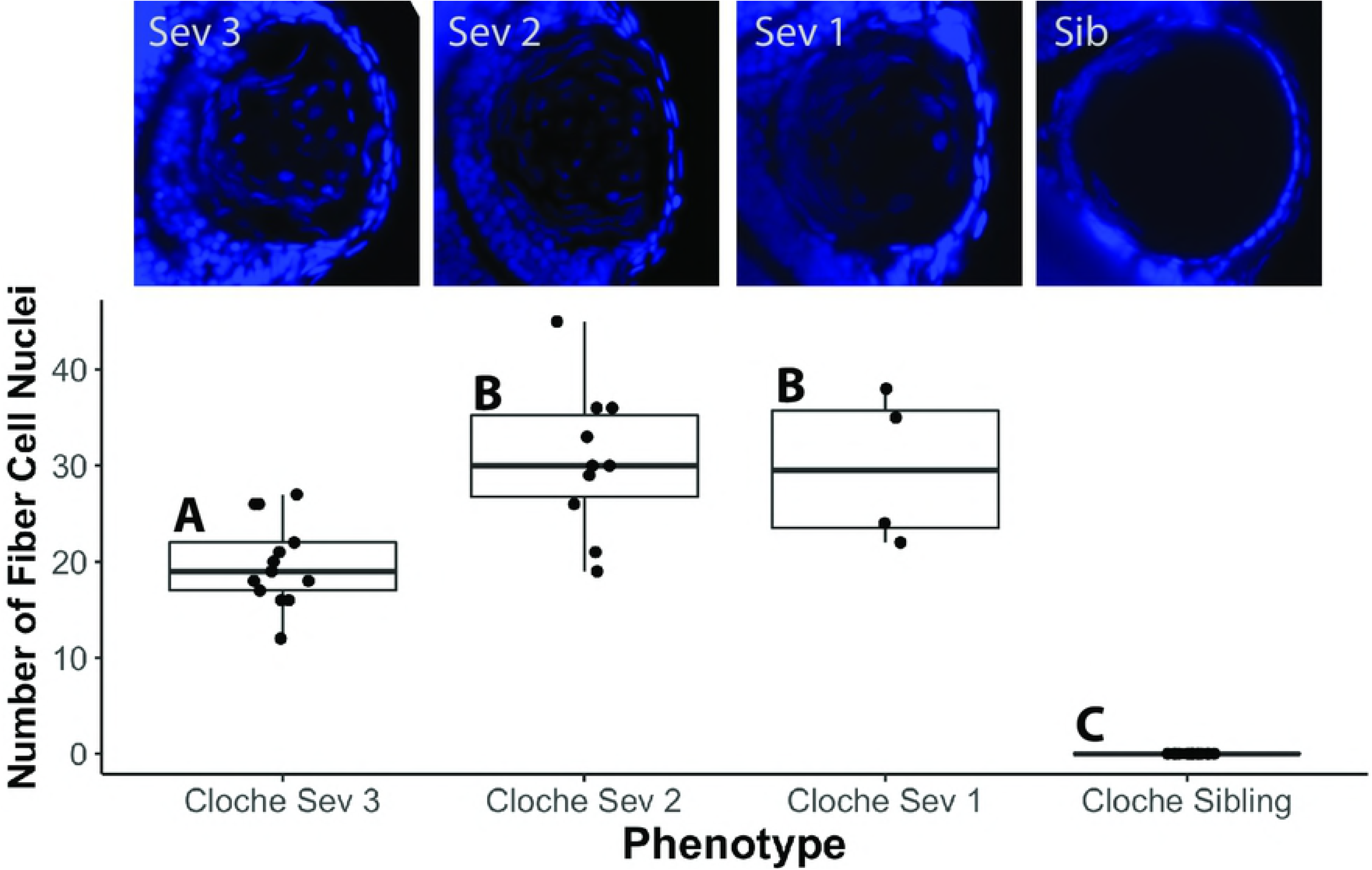
Quantification of retained fiber cell nuclei in *cloche* lenses of different phenotype severity compared to non-phenotype siblings by DAPI staining. Images above the graph show representative lenses for each severity type at 4 dpf. Fiber cell nuclei were significantly more abundant in all *cloche* lenses compared to siblings. Within cloche embryos, severity type 3 lenses (the most severe) contained fewer nuclei than severity type 2 or 1 (ANOVA *p* value < 0.0001; letters indicate statistical groups determined by Tukey Honest Significant Difference (HSD) post test).

### Few lens specific genes showed changes in expression in *cloche* embryos compared to wildtype

Goishi et al. [15] used semi-quantitative RT-PCR to show a decrease in αA-crystallin mRNA in 2, 3 and 4 dpf *cloche* embryos compared to wildtype embryos. We compared the expression of all three zebrafish α-crystallin genes (*cryaa, cryaba, cryabb*) at 4 dpf in *cloche* phenotype, *cloche* siblings and wildtype fishes by qRT-PCR and found no significant differences in expression for any of these genes (Fig 4).

**Figure 4.**
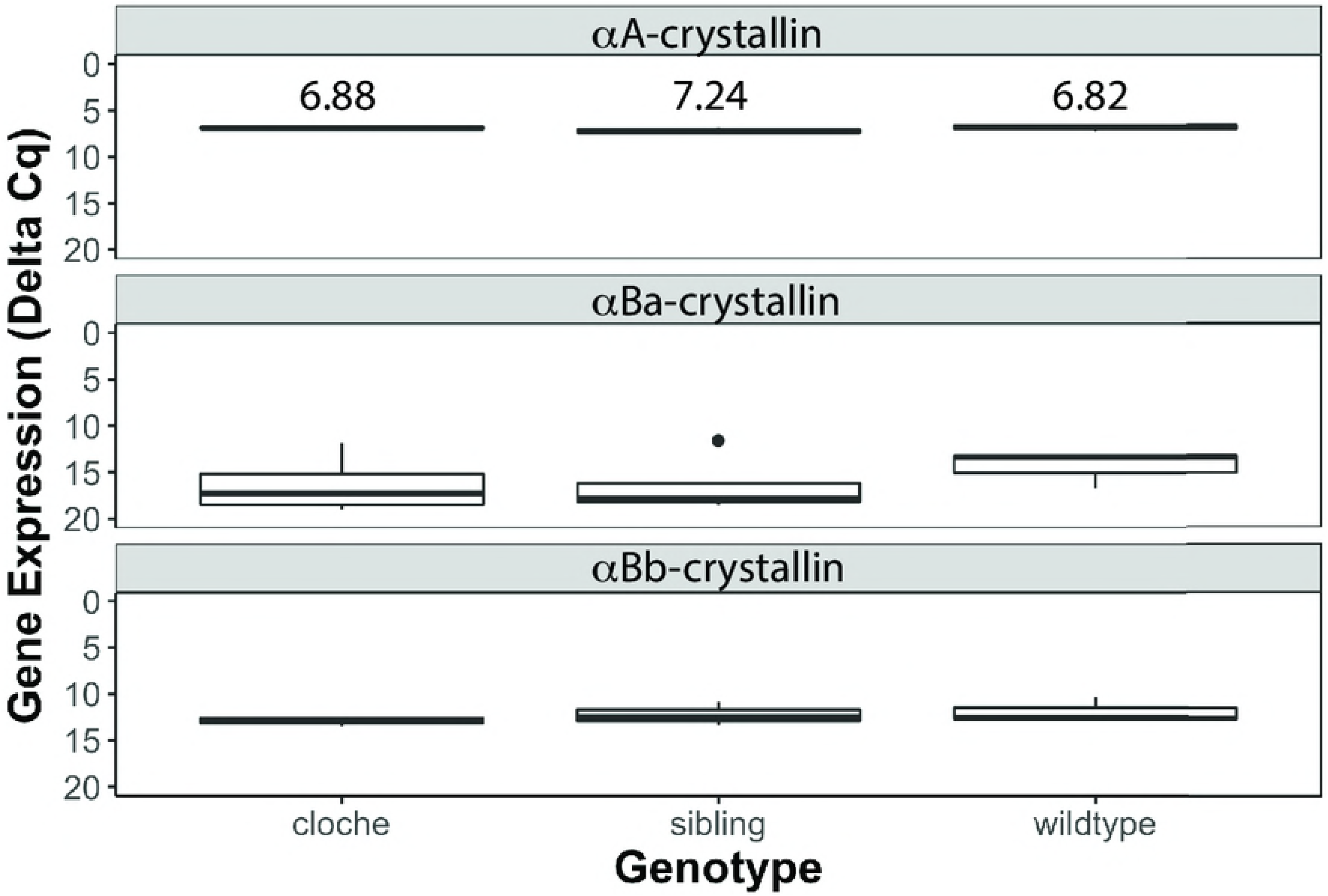
Quantitative RT-PCR analysis of α-crystallin expression at 4 dpf. mRNA levels for each of the three zebrafish α-crystallin genes were similar between *cloche* embryos, *cloche* siblings and wildtype embryos. αB-crystallin gene expression was low and measurements more variable at these stages, similar to what we have previously reported [5]. There were no statistical differences in delta Cq values for each gene between sample type (ANOVA; αA(F=1.6941, *p* value=0.261), αBa(F=0.491, *p* value=0.6293), αBb(F=0.6327, *p* value=0.5632). Each Cq value represents a biological triplicate for each sample normalized to two reference genes, with lower values indicating higher levels of gene expression. Delta Cq values are indicated for the αA-crystallin analysis.

We used RNA-Seq to analyze global gene expression differences between *cloche* phenotype embryos and wildtype embryos at 4 dpf. Our analysis collected over 400 million reads and identified transcripts from 25,281 genes, 1,329 of which were differentially expressed between *cloche* and wildtype embryos (Fig 5, Supplement 1). None of the three zebrafish α-crystallin genes were found to be statistically significantly differentially expressed in *cloche* embryos compared to wildtype. Very low expression was observed for *cryaba* and *cryabb*, consistent with our qRT-PCR data and publicly available developmental RNA-Seq data for wildtype zebrafish (https://www.ebi.ac.uk/gxa/experiments/E-ERAD-475). We did find genes for one α-crystallin and seven γm-crystallins that were downregulated in *cloche* embryos and one crystallin gene (*crybg1b*) that was upregulated (Fig 5; Supplement 2). Several transcription factors known to regulate lens development, such as *pax6a, foxe3, hsf4*, and *prox1a* were not differentially expressed in *cloche* embryos compared to wildtypes. However, *pax6b*, which is involved in cornea and lens development [33], was downregulated in *cloche*. The lens membrane protein gene *mipb* [31,34], which has been linked to cataract development, was also downregulated. The gene *sil1*, which has been linked to Marinesco-Sjögren syndrome (MSS) including lens cataract [35], was upregulated in *cloche* embryos while *fabp11*, a fatty acid binding gene linked to eye development [36], was downregulated.

**Figure 5.**
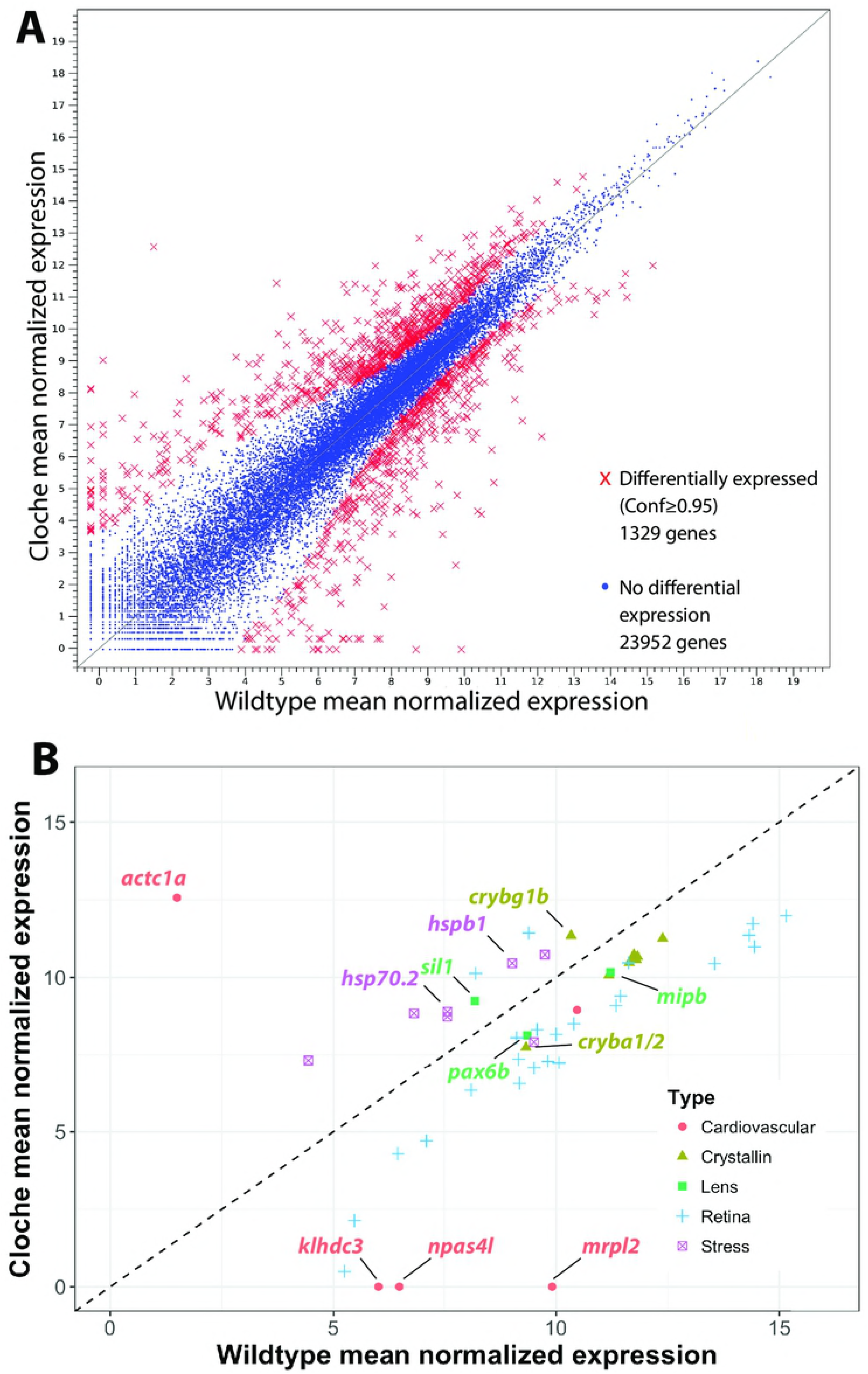
RNA-Seq identified 1,329 genes with differential expression between 4 dpf wildtype and *cloche* embryos. Transcripts were read from a total of 25,281 genes that are plotted by normalized expression levels in wildtype and *cloche* embryos (A). Panel B shows a subset of genes for lens crystallins, other lens related proteins, retinal related proteins and stress proteins. The two α-crystallin genes identified in the RNA-Seq analysis did not differ in expression between wildtype and cloche but are included for reference. Confidence values for determining differential expression were produced by bROC analysis.

Over 20 retina related genes were downregulated in *cloche* embryos and at least two (*hbegfa* and *odc1*) showed increased expression (Fig 5; Supplement 2). This overall reduction in retina related gene expression may reflect the reduced retinal cell proliferation, cell survival and differentiation of retinal cell types identified in this mutant [32].

Our RNA-Seq analysis successfully identified the expected loss of *klhdc3, mrpl2* and *npas4l* expression in *cloche* embryos, three genes identified as being lost in this mutant line [14]. Other genes involved in hematopoiesis (*lclat1*) and oxygen delivery (*hbae1.1, hbbe1.3, hbbe1.1*) were strongly down regulated in *cloche* embryos as well (Supplement 1). The expression of an anoxia responsive gene, *phd3*, was not significantly changed in *cloche* embryos, similar to findings at 3 dpf by Dhakal et al. [32] indicating that loss of blood circulation is not putting *cloche* embryos in anoxic stress. Interestingly, a number of heat shock protein genes (eg. *hspa13, hspb9, hspb1, hsp70.2, hsp70.3*) were upregulated in *cloche* embryos while one, *hspa41*, was downregulated (Fig 5; Supplement 2). Two genes identified through automated annotation of the zebrafish genome, both located on chromosome 15, were upregulated from essentially no detectable expression in wildtype to strong expression in *cloche* (*si:ch211-181d7.1_2* and *si:ch211-181d7.3*; Supplement 1). Both resulting proteins are predicted to bind ATP, but biological functions are not known.

### Characterization of native aA-crystallin promoter activity in *cloche* and wildtype lens

We compared the activity of two lens crystallin promoters in *cloche* embryos to test a previously published hypothesis that the native zebrafish α-crystallin promoter is downregulated in this mutant [15]. We also wanted to determine if one of these promoters would allow us to efficiently drive the expression of introduced proteins into *cloche* embryo lenses for future tests of their ability to suppress cataract formation. A native zebrafish αA-crystallin promoter (-1000/+1) produced lens GFP expression in similarly high percentages of 4 dpf *cloche, cloche-sibling* and wildtype embryos (Fig 6). The human βB1-crystallin promoter (-223/+61), however, produced lens GFP expression in a statistically significantly lower proportion of both *cloche* and *cloche*-siblings compared to the zebrafish αA-crystallin promoter (Yates Corrected *X^2^* test: *X^2^*=16.85, *p* value<0.001; *X^2^*=21.38, *p* value<0.001 respectively). There was no statistical difference in wildtype embryos (*X^2^*=3.35, *p* value>0.05). The visible intensity of GFP expression was statistically significantly higher with the αA-crystallin promoter in all three embryo types (t-test: *p* values = 1.72 X 10^-7^, 6.33 X 10^-6^ and 0.037 respectively).

**Figure 6.**
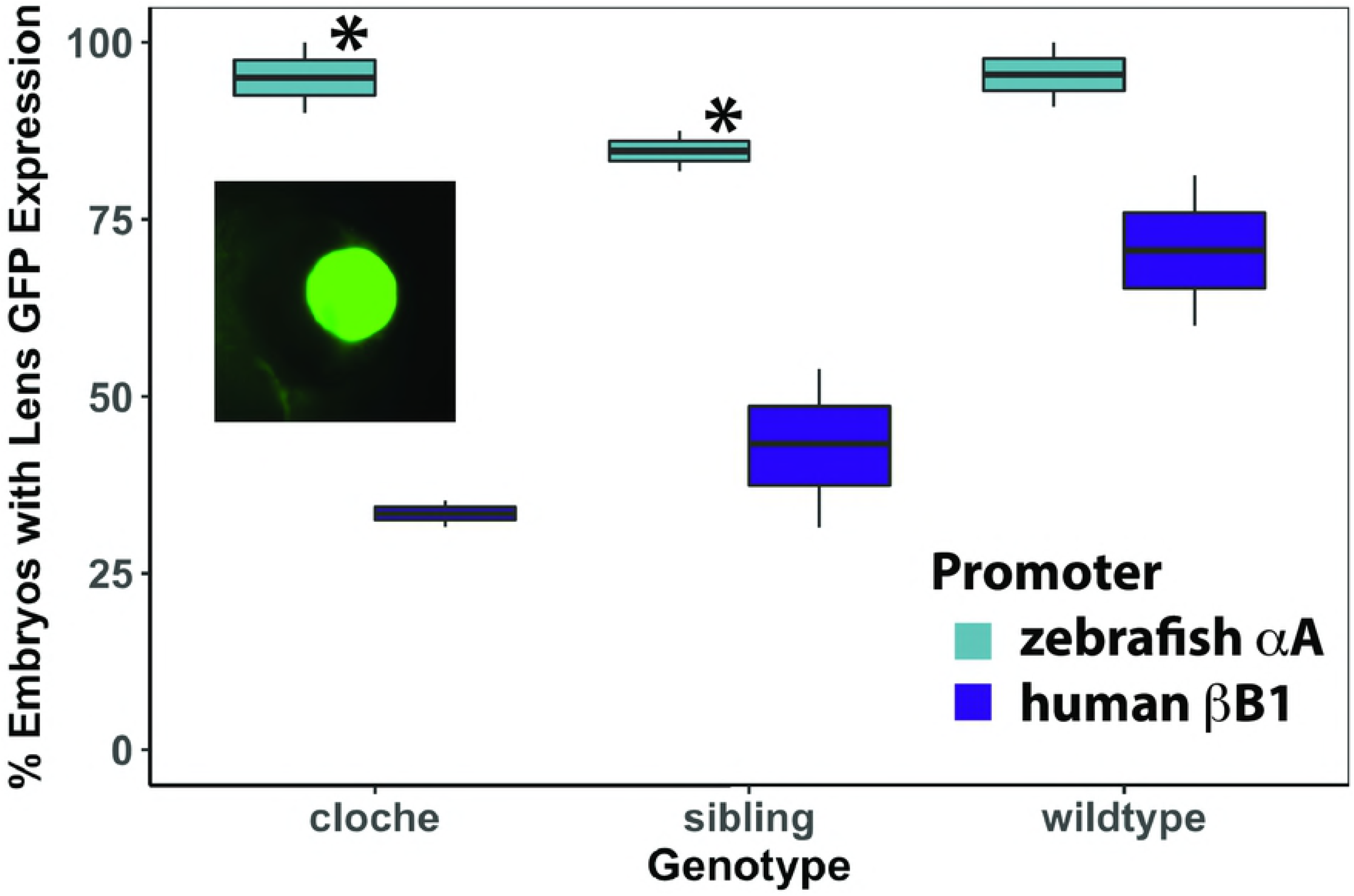
Percent of embryos with lens GFP expression after injection of zebrafish αA-crystallin promoter/GFP and human βB1-crystallin promoter/GFP plasmids. Data show that the native αA-crystallin promoter drives greater GFP expression in lens compared to the human bB1 promoter in *cloche* and non-phenotype siblings (Yates Corrected *X^2^* test: *X^2^*=16.85, *p* value<0.001; *X^2^* =21.38, *p* value<0.001 respectively), but this difference was not statistically significant in wildtype embryos (*X^2^* =3.35, *p* value>0.05). Each box and whisker blot represents two independent experiments (except for the βB1 sibling value which included three independent experiments). Each independent experiment included between 3 and 54 embryos at 4 dpf (median = 19). Inset shows an example of GFP lens expression in a *cloche* embryo produced by the αA-crystallin promoter.

A time series experiment provided additional data to show that the zebrafish αA-crystallin promoter is active in the lens of a higher proportion of embryos compared to the human bB1-crystallin promoter (Fig 7B). Interestingly, the human βB1-promoter did drive strong GFP expression in skeletal muscle and was active earlier than the αA-promoter (Fig 7B). These data combined embryos of all genotypes.

**Figure 7.**
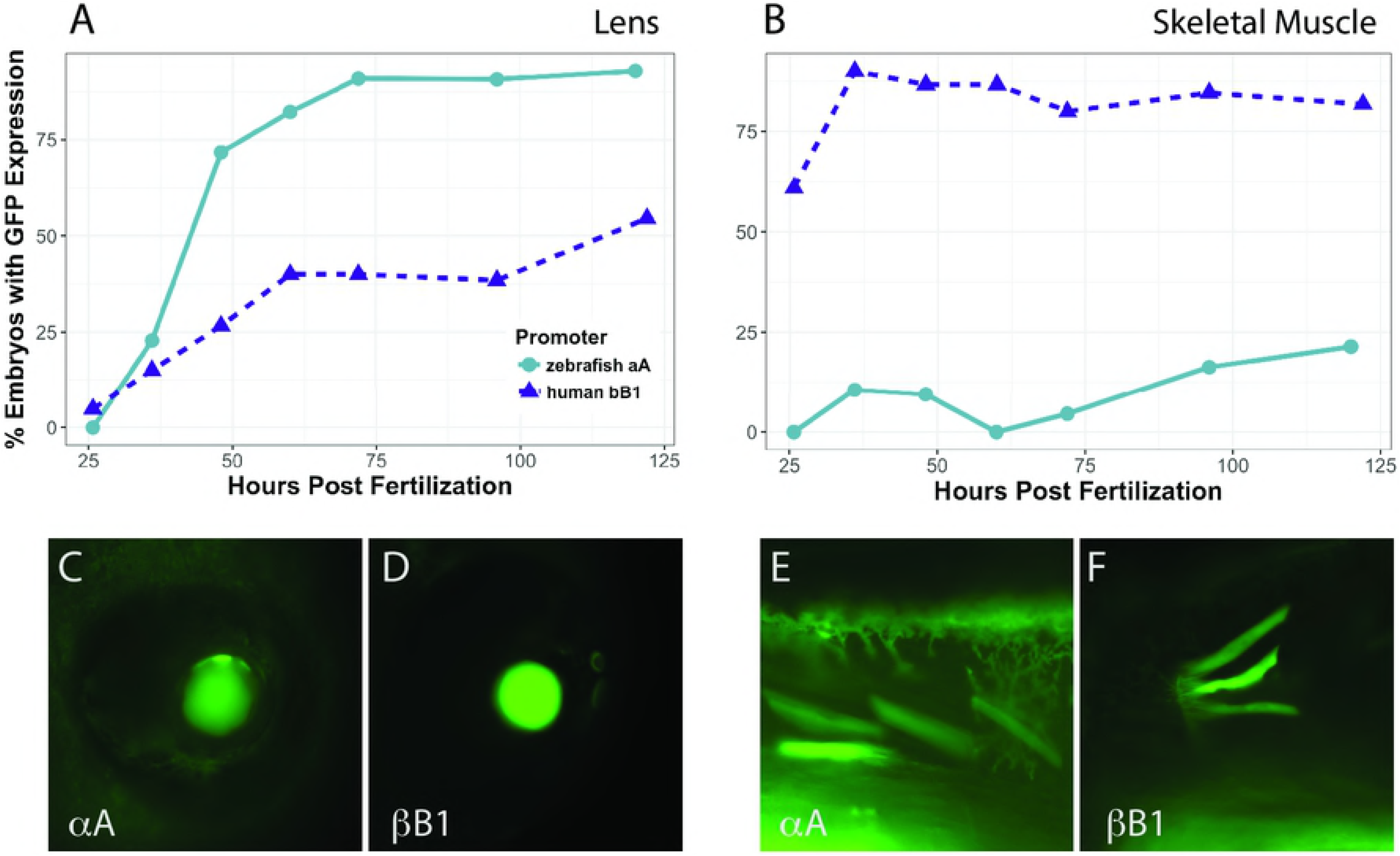
Timecourse of promoter activity in lens (A) and skeletal muscle (B) in all combined embryos. Zebrafish αA-crystallin promoter (circles) produced lens expression in a larger proportion of embryos by 36 hpf and drove GFP expression in over 85% of embryos by 72 hpf. The human βB1-crystallin promoter (triangles) drove surprisingly abundant expression in zebrafish skeletal muscle, but was less active in lens. Between 11 and 66 embryos were observed for each timepoint. Images C-F show representative examples of GFP expression with each promoter as indicated in lens (C and D) and skeletal muscle (E and F).

## Discussion

Our work and that from others has identified at least three features of the *cloche* lens phenotype. The lens is smaller than normal, shows arrested denucleation of fiber cells and contains a noticeable central irregularity that previous studies have shown scatters light [15,32]. Data in this present study show that lens size and retention of fiber cells are similar in all *cloche* lenses, while the central irregularity can vary in severity. Goishi et al. [15] showed that *cloche* lenses include aggregated γ-crystallins, which likely contributes to the visual roughness seen by DIC microscopy. The variability in appearance of this lens irregularity suggests that it is stochastic. Whatever physiological and/or molecular mechanisms lead to reduced eye size and fiber cell denucleation arrest do not similarly dictate cataract formation, but may make the lens more prone to protein aggregation.

The *cloche* phenotype embryo does not appear to transcribe significantly reduced levels of αA-crystallin mRNA compared to its non-phenotype siblings or wildtype zebrafish at 4 days post fertilization. This conclusion is supported by qRT-PCR and RNA-Seq data, as well as promoter expression data that show similar levels of GFP expression in *cloche* embryos and wildtype when driven by a native zebrafish αA-crystallin promoter. Together, these data suggest that the *cloche* lens cataract is not triggered by a reduction in αA-crystallin expression. There are possible reasons why this result differs from the decrease in αA-crystallin expression reported by Goishi et al. [15]. First, while both studies used the same *cloche* allele (*m39*) we are likely propagating that allele in different genetic backgrounds. These genetic differences could influence *cryaa* expression. Second, we have used different methods to measure *cryaa* expression (qRT-PCR and RNA-Seq compared to end point RT-PCR). It is worth noting that our RNA-Seq data showed lower *cryaa* mean normalized expression in *cloche* embryos compared to wildtype (-0.85 fold, bROC confidence=0.937) but this difference was not statistically significant. While we cannot exclude the possibility that there is some reduction in *cryaa* expression in *cloche* embryos, we believe that the data in this study provide strong evidence that loss of αA-crystallin is not the cause of the *cloche* cataract.

Previously published knockout studies also suggest that loss of αA-crystallin would not produce the severity of lens phenotype seen in the *cloche* mutant. The well characterized *cryaa* knockout mouse does not exhibit a lens cataract until seven weeks of age [37], later than the comparable *cloche* developmental stage examined here and in past studies. A recently published zebrafish TALEN knockout study identified a lens phenotype after disabling *cryaa* [18]. However, the reported lens phenotype is more subtle than found in the *cloche* lens. Expression of introduced αA-crystallin can reduce light scatter in the *cloche* lens, indicating rescue of the phenotype by addition of this protein [15,18]. But a reasonable alternate hypothesis is that introduction of additional chaperoning protein may reduce protein aggregation no matter what its initial cause. Based on the data in this present study and findings from previously published work we propose that reduction in αA-crystallin protein is not the sole cause of the *cloche* lens phenotype and suggest that another mechanism initiates the lens defects.

While the ultimate cause of the *cloche* lens cataract remains unknown, our RNA-Seq data suggest that dysregulation of known lens development-related transcription factors is not involved. The majority of the lens associated genes that do differ in expression between *cloche* phenotype and wildtype embryos code for structural proteins and are downregulated. The same pattern is seen in retinal genes, possibly reflecting reduced eye size caused by inhibition of ocular tissue development and growth [32]. Interestingly, the opposite may be occurring with heart size as an increased number of cardiomyocytes in the *cloche* mutant could be driving increased expression of *actc1a* [38,39]. The increase in expression of multiple stress response genes for both larger Hsp70 proteins and smaller Hspb proteins suggests a general physiological stress response in the *cloche* embryo triggered by the lack of hematopoiesis. Our RNA-Seq results are consistent with an hypothesis that *cloche* cataract formation is triggered by a general physiological stress and not a specific error in lens development gene regulatory networks. While cataract formation induced by this stress may be variable, leading to a range of phenotype severity, the reduction in *cloche* embryo lens diameter and delay in fiber cell differentiation is a more uniform response. It is possible that abnormal lens regulatory signaling in the *cloche* mutant is transient, and that our RNA-Seq analysis on 4 dpf embryos may have missed it. Resolving this possibility will require a future RNA-Seq timeseries analysis that covers key stages in lens development (lens placode delamination at 16 hpf; first fiber cell differentiation at 30 hpf; initial fiber cell denucleation at 72 hpf; [40]). For now we can conclude that the *cloche* lens phenotype, including the persistent loss of fiber cell denucleation, does not result from a noticeable change in the use of known lens gene regulatory networks.

Our comparison of the ability of two crystallin promoters to drive GFP expression in the *cloche* lens is an important step towards using the *cloche* cataract as a model for protein aggregation and cataract prevention. Based on our results a native zebrafish promoter containing 1 kb of upstream sequence from the start codon can effectively drive protein expression almost exclusively in the lens and should be a good tool for transiently expressing introduced proteins to test their impact on cataract formation. These future studies should include proteomic analysis to quantify levels of introduced protein expression. Our unexpected finding that the human βB1-crystallin promoter drove lower GFP expression in *cloche* embryos compared to wildtype serves as a cautionary note that promoter activity needs to be assessed before they are used in experiments testing the effects of introduced lens proteins.

The lack of noticeable changes to known molecular mechanisms underlying lens development in *cloche* embryos suggests that the presence of cataract in this mutant might not result from disturbances in lens specific gene regulation. Instead, the *cloche* lens may simply be responding to a general physiological stress, and changes to the expression of some lens crystallins may be a byproduct of, and not directly related to, a disturbance in crystallin-specific gene regulatory networks. However, additional time points added to our RNA-Seq analysis may provide an opportunity to observe transient changes in lens gene regulatory networks or uncover novel signaling molecules that contribute to lens development. What seems clear is that the *cloche* lens phenotype does not result from a significant loss of αA-crystallin expression. Whether or not the *cloche* zebrafish can provide further insights into the regulation of lens development, the presence of its lens cataract can be used to study lens crystallin aggregation and cataract prevention.

## Acknowledgements

This work was supported by an R15 AREA grant from the National Eye Institute (EY013535) to MP and from grants to support faculty/student research from the Provost Office of Ashland University and a summer student research stipend was provided as part of a Choose Ohio First scholarship grant to Ashland University to KLM. The *cloche^m39^* line was provided by the Zon laboratory at Harvard University and the human βB1-crystallin promoter construct was provided by the Hall lab at UC Irvine. Emmaline Kepp helped with analysis of the *cloche* phenotype. We would like to thank the Busch-Nentwich lab for providing publically available developmental zebrafish time series RNA-seq data used to validate the novel RNA-Seq data presented in this paper. Kirsten Lampi, Elizabeth LeClair and Saulius Sumanas provided feedback on the manuscript.

## Supplement Captions

**Supplemental Table 1.** List of 50 genes with the greatest bROC confidence levels for change in expression at 4 dpf. Gene symbol, mean normalized expression values for wildtype and cloche, fold change, confidence level and gene ontology (GO) biological or molecular processes are shown. Some GO terms have not been identified and are left blank. Fold changes calculated by the bootstrapping bROC method are smaller and more conservative than those determined by non-bootstrapping methods.

**Supplemental Table 2.** A subset of differentially expressed genes that are plotted in Fig 5B. Normalized expression was averaged across the three biological replicates for wildtype and *cloche* embryos. Fold change, bROC confidence levels and general biological function are indicated. Fold changes calculated by the bootstrapping bROC method are smaller and more conservative than those determined by non-bootstrapping methods.

